# Beyond species: why ecological interaction networks vary through space and time

**DOI:** 10.1101/001677

**Authors:** T. Poisot, D.B. Stouffer, D. Gravel

## Abstract

Community ecology is tasked with the considerable challenge of predicting the structure, and properties, of emerging ecosystems. It requires the ability to understand how and why species interact, as this will allow the development of mechanism-based predictive models, and as such to better characterize how ecological mechanisms act locally on the existence of interspecific interactions. Here we argue that the current conceptualization of species interaction networks is ill-suited for this task. Instead, we propose that future research must start to account for the intrinsic variability of species interactions, then scale up from here onto complex networks. This can be accomplished simply by recognizing that there exists intra-specific variability, in traits or properties related to the establishment of species interactions. By shifting the scale towards population-based processes, we show that this new approach will improve our predictive ability and mechanistic understanding of how species interact over large spatial or temporal scales.

## Introduction

Interactions between species are the driving force behind ecological dynamics within communities (Berlow et al. 2009). Likely for this reason more than any, the structure of communities have been described by species interaction networks for over a century (Dunne 2006). Formally an ecological network is a mathematical and conceptual representation of both *species*, and the *interactions* they establish. Behind this conceptual framework is a rich and expanding literature whose primary focus has been to quantify how numerical and statistical properties of networks relate to their robustness (Dunne et al. 2002), productivity (Duffy et al. 2007), or tolerance to extinction (Memmott et al. 2004). Although this approach classically focused on food webs (Ings et al. 2009), it has proved particularly successful because it can be applied equally to all types of ecological interactions (Kéfi et al. 2012).

This body of literature generally assumes that, short of changes in local densities due to ecological dynamics, networks are inherently *static* objects. This assumption calls into question the relevance of network studies at biogeographic scales. More explicitly, if two species are known to interact at one location, it is often assumed that they will interact whenever and wherever they co-occur (see *e.g.* Havens 1992); this neglects the fact that local environmental conditions, species states, and community composition can intervene in the realization of interactions. More recently, however, it has been established that networks are *dynamic* objects that have structured variation in *α*, *β*, and *γ* diversity, not only with regard to the change of species composition at different locations but also to the fact that the same species will interact in different ways over time or across their area of co-occurrence (Poisot et al. 2012). Of these sources of variation in networks, the change of species composition has been addressed explicitly in the context of networks (Gravel et al. 2011, Dáttilo et al. 2013) and within classical meta-community theory. However, because this literature still tends to assume that interactions happen consistently between species wherever they co-occur, it is ill-suited to address network variation as a whole and needs be supplemented with new concepts and mechanisms. Within the current paradigm, interactions are established between species and are an immutable “property” of a species pair. Starting from empirical observations, expert knowledge, or literature surveys, one could collect a list of interactions for any given species pool. Several studies used this approach to extrapolate the structure of networks over time and space (Havens 1992, Piechnik et al. 2008, Baiser et al. 2012) by considering that the network at *any* location is composed of *all* of the potential interactions known for this species pool. This stands in stark contrast with recent results showing that (i) the identities of interacting species vary over space and (ii) the dissimilarity of interactions is not related to the dissimilarity in species composition (Poisot et al. 2012). The current conceptual and operational tools to study networks therefore leaves us poorly equipped to understand the causes of this variation. In this paper, we propose to shift the research agenda towards understanding the mechanisms involved in the variability of co-occurring species interactions.

In contrast to the current paradigm, we propose that future research on interaction networks should be guided by the following principles: the existence of an interaction between two species is the result of a stochastic process involving (i) local traits distributions, (ii) local abundances, and (iii) higher-order effects by the local environment or species acting “at a distance” on the interaction; regionally, the observation of interactions results of the accumulation of local observations. This approach is outlined in Box 1. Although this proposal is a radical yet intuitive change in the way we think about ecological network structure, we demonstrate in this paper that it is well supported by empirical and theoretical results alike. Furthermore, our new perspective is well placed to open the door to novel predictive approaches integrating a range of key ecological mechanisms. Notably, we propose in Box 2 that this approach facilitates the study of indirect interactions, for which predictive approaches have long proved elusive (Tack et al. 2011).

#### Box 1 A mathematical framework for population-level interactions

We propose that the occurrence (and intensity) of ecological interactions at the population level relies on several factors, including relative local abundances and local trait distributions. It is important to tease apart these different factors so as to better disentangle neutral and niche processes. We propose that these different effects can adequately be partitioned using the model

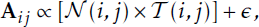

where *𝒩* is a function giving the probability that species *i* and *j* interact *based only on their local abundances* (that is, the probability of encounter), and *𝒯* is a function giving the *per encounter* probability that species *i* and *j* interact *based on their trait values*. The term *ε;* accounts for all higher-order effects, such as indirect interactions, local impact of environmental conditions on the interaction, and impact of co-occurring species. Both of these functions can take any form needed. In several papers, *𝒩*(*i, j*) was expressed as n_*i*_ × n_*j*_, where n is a vector of relative abundances (Canard et al. 2014). The expression of *𝒯* can in most cases be derived from mechanistic hypotheses about the observation. For example, Gravel et al. (2013) used the niche model of Williams and Martinez (2000) to predict interactions with the simple rule that *𝒯* (*i*, *j*) = 1 if *i* can consume *j* based on allometric rules, and 0 otherwise. Following Rohr et al. (2010), the expression of *𝒯* can be based on latent variables rather than actual trait values. This simple formulation could be used to partition, at the level of individual interactions, the relative importance of density-dependent and trait-based processes using variance decomposition. Most importantly, it predicts (i) how each of these components will vary over space and (ii) how the structure of the network will be affected by, for example, changes in local abundances or trait distributions. The results provided by this framework will only be as good as the empirical data used, and there is a dire need for a methodological discussion about how “predictor” variables (traits, population sizes, etc.) should be measured in the field, in a way that is not biased by the observation of the interactions. This will prove challenging for some types of interactions; *e.g.* estimating the population size of parasites is often contingent upon catching and examining hosts. Understanding non-independence between these variables in a system-specific way is a crucial point.

This model can further be extended in a spatial context, as

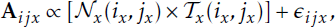

in which *i*_*x*_ is the population of species *i* at site *x*. In this formulation, the *ǫ* term could include the spatial variation of interaction between *i* and *j* over sites, and the covariance between the observed presence of this interaction and the occurrence of species *i* and *j*. This can, for example, help address situations in which the selection of prey items is determined by traits, but also by behavioral choices. Most importantly, this model differs from the previous one in that each site *x* is characterized by a set of functions *𝒩_x_*, *𝒯_x_* that may not be identical for all sites considered. For example, the same predator may prefer different prey items in different locations, which will require the use of a different form for *𝒯* across the range of locations. Gravel et al. (2013) show that it is possible to derive robust approximation for the *𝒯* function even with incomplete set of data, which gives hope that this framework can be applied even when all species information is not known at all sites (which would be an unrealistic requirement for most realistic systems). Both of these models can be used to partition the variance from existing data or to test which trait-matching function best describes the observed interactions. They also provide a solid platform for dynamical simulations in that they will allow re-wiring the interaction network as a function of trait change and to generate simulations that are explicit about the variability of interactions.

#### Box 2 Population-level interactions in the classical modelling framework

As noted in the main text, most studies of ecological networks—particularly food webs—regard the adjacency matrix A as a fixed entity that specifies observable interactions on the basis of whether two species co-occur or not. Given this assumption, there is a lengthy history of trying to understand how the strength or organization of these interactions influence the dynamic behavior of species abundance (May 1973). Often, such models take the form

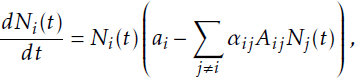

where *a*_*i*_ is the growth rate of species *i* (and could, in principle, depend on other species’ abundances *N)* and *α*_*ij*_ is the strength of the effect of *j* on *i*. In this or just about any related model, direct species-species interaction can influence species abundances but their abundances *never* feedback and influence the *per capita* interaction coefficients *α*_*ij*_. They do, however, affect the realized interactions, which are defined by *α*_*ij*_ *𝒩*_*i*_ (*𝒯*)*𝒩*_*j*_ (*𝒯*), something which is also the case when considering more complicated functional responses (Koen-Alonso 2007).

More recently, there have been multiple attempts to approach the problem from the other side. Namely, to understand how factors such as species’ abundance and/or trait distributions influence the occurrence of the interactions themselves (Box 1). One potential drawback to that approach, however, is that it still adopts the assumption that the observation of any interaction *A*_*ij*_ is only an explicit function of the properties of species *i* and *j* (traits and co-occurrence).

Since dynamic models demonstrate quite clearly that non-interacting species can alter each others’ abundances (*e.g.* via apparent competition (Holt and Kotler 1987)), this is a deeply-ingrained inconsistency between the two approaches. Such a simplification does increase the analytical tractability of the problem (Allesina and Tang 2012), but there is little, if any, guarantee that it is ecologically accurate. In our opinion, the “higher-effects” term *є* in the models presented in Box 1 is the one with the least straightforward expectations, but it may also prove to be the most important if we wish to accurately describe all of these indirect effects.

A similar problem actually arises in the typical statistical framework for predicting interaction occurrence. Often, one attempts to “decompose” interactions into the component that is explained by species’ abundances and the component explained by species’ traits (e.g., Box 1). Just like how the underlying functions *𝒩* and *𝒯* could vary across sites, there could also be feedback between species’ abundances and traits, in the same way that we have outlined the feedback between interactions and species’ abundances. In fact, given the increasing evidence for the evolutionary role of species-species interactions in explaining extant biodiversity and their underlying traits (Janzen and Martin 1982, Herrera et al. 2002), a framework which assumes relative independence of these different phenomenon is likely starting from an overly-simplified perspective.

Since the next generation of predictive biogeographic models will need to account for species interactions (Thuiller et al. 2013), it is crucial not to underestimate the fact that they are intrinsically variable and exhibit a geographic variability of their own. Indeed, investigating the impact of species interactions on species distributions only makes sense under the implicit assumption that species interactions themselves vary over biogeographical scales. Models of species distributions will therefore increase their predictive ability if they account for the variability of ecological interactions. In turn, tighter coupling between species-distribution and interaction-distribution models will provide mode accurate predictions of the properties of emerging ecosystems (Gilman et al. 2010, Estes et al. 2011) and the spatial variability of properties between existing ecosystems. By paying more attention to the variability of species interactions, the field of biogeography will be able to re-visit classical observations typically explained by species-level mechanisms; for example, how does community complexity and function vary along latitudinal gradients, is there information hidden in the co-occurrence or avoidance of species interactions, etc. This predictive effort is made all the more important as both the phenology (Parmesan 2007) and ranges (Devictor et al. 2012) of species occupying different positions in their interactions networks are affected differently by climate change. Predicting that species will move and change while interactions remain the same is probably a very conservative approach to estimating the changes to come, and building explicitly on biological mechanisms is one possible way to overcome this limitation.

In this paper, we outline the mechanisms that are involved in the variability of species interactions over time, space, and environmental gradients. We discuss how they will affect the structure of ecological networks, and how these mechanisms can be integrated into new predictive and statistical models (Box 1). Most importantly, we show that this approach integrates classical community ecology thinking and biogeographic questions (Box 2) and will ultimately result in a better understanding of the structure of ecological communities.

## The dynamic nature of ecological interaction networks

Recent studies on the sensitivity of network structure to environmental change provide some context for the study of dynamic networks. Menke et al. (2012) showed that the structure of a plant–frugivore network changed along a forest–farmland gradient. At the edges between two habitats, species were on average less specialized and interacted more evenly with a larger number of partners than they did in habitat cores. Differences in network structure have also been observed within forest strata that differ in their proximity to the canopy and visitation by birds (Schleuning et al. 2011). Tylianakis et al. (2007) reports a *stronger* signal of spatial interaction turnover when working with quantitative rather than binary interactions, highlighting the importance of *measuring* rather than assuming (or simply reporting) the existence of interactions. Eveleigh et al. (2007) demonstrated that outbreaks of the spruce budworm were associated with changes in the structure of its trophic network, both in terms of species observed and their interactions. Poisot et al. (2011) used a microbial system of hosts and pathogens to study the impact of productivity gradients on realized infection; when the species were moved from high to medium to low productivity, some interactions were lost and others were gained. As a whole, these results suggest that the existence, and properties, of an interaction are not only contingent on the presence of the two species involved but may also require particular environmental conditions, including the presence or absence of species not directly involved in the interaction.

We argue here that there are three broadly-defined classes of mechanisms that ultimately determine the realization of species interactions. First, species must be locally abundant enough for their individuals to meet; this is the so-called “neutral” perspective of interactions. Second, there must be phenological or trait matching between individuals, such that an interaction will actually occur given that the encounter takes place. Finally, the realization of an interaction is regulated by the interacting organisms’ surroundings and should be studied in the context of indirect interactions.

## Population dynamics and neutral processes

Over the recent years, the concept of neutral dynamics has left a clear imprint on the analysis of ecological network structure, most notably in bipartite networks (Blüthgen et al. 2006). Re-analysis of several host–parasite datasets, for example, showed that changes in local species abundances triggers variation in parasite specificity (Vazquez et al. 2005). More generally, it is possible to predict the structure of trophic interactions (Canard et al. 2012) and host-parasite communities (Canard et al. 2014) given only minimal assumptions about the distribution of species abundance. In this section, we review recent studies investigating the consequences of neutral dynamics on the structure of interaction networks and show how variations in population size can lead directly to interaction turnover.

### The basic processes

As noted previously, for an interaction to occur between individuals from two populations, these individuals must first meet, then interact. Assuming that two populations occupy the same location and are active at the same time of the day/year, then the likelihood of an interaction is roughly proportional to the product of their relative abundance (Vázquez et al. 2007). This means that individuals from two large populations are more likely to interact than individuals from two small populations, simply because they tend to meet more often. This approach can also be extended to the prediction of interaction strength (Blüthgen et al. 2006, Vázquez et al. 2007), *i.e.* how strong the consequences of the interaction will be. The neutral perspective predicts that locally-abundant species should have more partners and that locally-rare species should appear more specialized. In a purely neutral model (*i.e.* interactions happen entirely by chance, although the determinants of abundance can still be non-neutral), the identities of species do not matter, and it becomes easy to understand how the structure of local networks can vary since species vary regionally in abundance. Canard et al. (2012) proposed the term of “neutrally forbidden links” to refer to interactions that are phenologically feasible but not realized because of the underlying population size distribution. The identity of these neutrally forbidden links will vary over time and space, either due to stochastic changes in population sizes or because population size responds deterministically (*i.e.* non-neutrally) to extrinsic drivers.

### Benefits for network analysis

It is important to understand how local variations in abundance, whether neutral or not, cascade up to affect the structure of interaction networks. One approach is to use simple statistical models to quantify the effect of population sizes on local interaction occurrence or strength (see *e.g.* Krishna et al. 2008). These models can be extended to remove the contribution of neutrality to link strength, allowing us to work directly on the interactions as they are determined by traits (Box 1). Doing so allows us to compare the variation of neutral and non-neutral components of network structure over space and time. To achieve this goal, however, it is essential that empirical interaction networks (i) are replicated and (ii) include independent measurements of population sizes.

An additional benefit of such sampling is that these data will also help refine neutral theory. Wootton (2005) made the point that deviations of empirical communities from neutral predictions were most often explained by species trophic interactions which are notoriously, albeit intentionally, absent from the original formulation of the theory (Hubbell 2001). Merging the two views will increase our explanatory power, and provide new ways to test neutral theory in interactive communities; it will also offer a new opportunity, namely to complete the integration of network structure with population dynamics. To date, most studies have focused on the effects of a species’ position within a food web on the dynamics of its biomass or abundance (Brose et al. 2006, Berlow et al. 2009, Stouffer et al. 2011, Saavedra et al. 2011). Adopting this neutral perspective brings things full circle since the abundance of a species will also dictate its position in the network: changes in abundance can lead to interactions being gained or lost, and these changes in abundance are in part caused by existing interactions (Box 2). For this reason, there is a potential to link species and interaction dynamics and, more importantly, to do so in a way which accounts for the interplay between the two. From a practical point of view, this requires repeated sampling of a system through time, so that changes in relative abundances can be related to changes in interaction strength (Yeakel et al. 2012). Importantly, embracing the neutral view will force us to reconsider the causal relationship between resource dynamics and interaction strength since, in a neutral context, both are necessarily interdependent.

## Traits matching in space and time

Once individuals meet, whether they will interact is widely thought to be the product of an array of behavioral, phenotypic, and cultural aspects that can conveniently be referred to as a “trait-based process”. Two populations can interact when their traits values allow it, *e.g.* viruses are able to overcome host resistance, predators can capture the preys, trees provide enough shading for shorter grasses to grow. Non-matching traits will effectively prevent the existence of an interaction, as demonstrated by Olesen et al. (2011). Under this perspective, the existence of interactions can be mapped onto trait values, and interaction networks will consequently vary along with variation in local trait distribution. In this section, we review how trait-based processes impact network structure, how they can create variation, and the perspective they open for an evolutionary approach.

### The basic processes

There is considerable evidence that, at the species level, interaction partners are selected on the grounds of matching trait values. Random networks built on these rules exhibit realistic structural properties (Williams and Martinez 2000, Stouffer et al. 2005). Trait values, however, vary from population to population within species; it is therefore expected that the local interactions will be contingent upon traits spatial distribution (Figure 2). The fact that a species’ niche can appear large if it is the aggregation of narrow but differentiated individual or population niches is now well established (Bolnick et al. 2003, Devictor et al. 2010a) and has also reinforced the need to understand intra-specific trait variation to describe the structure and dynamics of communities (Woodward et al. 2010, Bolnick et al. 2011). Nevertheless, this notion has yet to percolate into the literature on network structure despite its most profound consequence: a species appearing generalist at the regional scale can easily be specialized in *each* of the patches it occupies. This reality has long been recognized by functional ecologists, which are now increasingly predicting the *variance* in traits of different populations within a species (Violle et al. 2012).

**Figure 1:**
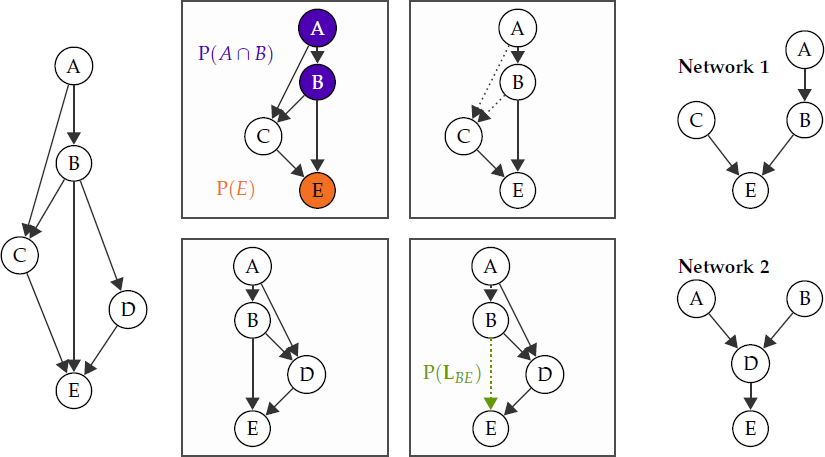
An illustration of the metaweb concept. In its simplest form, a metaweb is the list of all possible species and interactions between them for the system being studied, at the regional level (far left side). Everything that is ultimately observed in nature is a *realisation* of the metaweb (far right side), *i.e.* the resulting network after several sorting processes have occurred (central panel). First, species and species pairs have different probabilities to be observed (top panels). Second, as a consequence of the mechanisms we outline in this paper, not all interactions have the same probability to occur at any given site (bottom panels, see Box 1).

**Figure 2:**
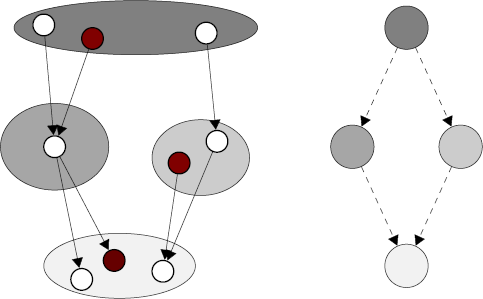
The left-hand side of this figure represents possible interactions between populations (circles) of four species (ellipses), and the aggregated species interaction network on the right. In this example, the populations and species level networks have divergent properties, and the inference on the system dynamics are likely to be different depending on the level of observation. More importantly, if the three populations highlighted in red were to co-occur, there would be no interactions between them, whereas the species-level network would predict a linear chain.

Empirically, there are several examples of intraspecific trait variation resulting in extreme interaction turnover. A particularly spectacular example was identified by Ohba (2011) who describes how a giant waterbug is able to get hold of, and eventually consume, juveniles from a turtle species. This interaction can only happen when the turtle is small enough for the morphotraits of the bug to allow it to consume the turtle, and as such will vary throughout the developmental cycle of both species. Choh et al. (2012) demonstrated through behavioral assays that prey which evaded predation when young were more likely to consume juvenile predators than the “naive” individuals; their past interactions shaped behavioral traits that alter the network structure over time. These examples show that trait-based effects on networks can be observed even in the absence of genotypic variation (although we discuss this in the next section).

From a trait-based perspective, the existence of an interaction is an emergent property of the trait distribution of local populations: variations in one or both of these distributions, regardless of the mechanism involved (development, selection, plasticity, environment), are likely to alter the interaction. Importantly, when interaction-driving traits are subject to environmental forcing (for example, body size is expected to be lower in warm environments, Angilletta et al. (2004)), there can be covariation between environmental conditions and the occurrence of interactions. Woodward et al. (2012) used macrocosms to experimentally demonstrate that changes in food-web structure happen at the same time as changes in species body mass distribution. Integrating trait variation over gradients will provide more predictive power to models of community response to environmental change.

### Benefits for network analysis

Linking spatial and temporal trait variation with network variation will help identify the mechanistic basis of network dissimilarity. From a sampling point of view, having enough data requires that, when interactions are recorded, they are coupled with trait measurements. Importantly, these measurements cannot merely be extracted from a reference database because interactions are driven by *local* trait values and their matching across populations from different species. Within our overarching statistical framework (Box 1), we expect that (i) network variability at the *regional* scale will be dependent on the variation of populations’ traits, and (ii) variation between any series of networks will depend on the *covariance* between species traits. Although it requires considerably larger quantities of data to test, this approach should allow us to infer *a priori* network variation. This next generation of data will also help link variation of network structure to variation of environmental conditions. Price (2003) shows how specific biomechanical responses to water input in shrubs can have pleiotropic effects on traits involved in the interaction with insects. In this system, the difference in network structure can be explained because (i) trait values determine the existence of an interaction, and (ii) environmental features determine trait values. We have little doubt that future empirical studies will provide similar mechanistic narratives.

At larger temporal scales, the current distribution of traits also reflects past evolutionary history (Diniz-Filho and Bini 2008). Recognizing this important fact offers an opportunity to approach the evolutionary dynamics and variation of networks. Correlations between different species’ traits, and between traits and fitness, drive coevolutionary dynamics (Gomulkiewicz et al. 2000, Nuismer et al. 2003). Both of these correlations vary over space and time (Thompson 2005), creating patchiness in the processes and outcomes of coevolution. Trait structure and trait correlations are also disrupted by migration (Gandon et al. 2008, Burdon and Thrall 2009). Ultimately, understanding of how ecological and evolutionary trait dynamics affect network structure will provide a mechanistic basis for the historical signal found in contemporary network structures (Rezende et al. 2007, Eklof et al. 2011, Baskerville et al. 2011, Stouffer et al. 2012).

## Beyond direct interactions

In this section, we argue that, although networks are built around observations of direct interactions like predation or pollination, they also offer a compelling tool with which to address indirect effects on the existence and strength of interactions. Any direct interaction arises from the “physical” interaction of only two species, and, as we have already detailed, these can be modified by local relative abundances and/or species traits. Indirect interactions, on the other hand, are established through the involvement of another party than the two focal species, either through cascading effects (herbivorous species compete with insect laying eggs on plants) or through physical mediation of the environment (bacterial exudates increase the bio-availability of iron for all bacterial species; plants with large foliage provide shade for smaller species). As we discuss in this section, the fact that many (if not all) interactions are indirectly affected by the presence of other species (i) has relevance for understanding the variation of interaction network structure and (ii) can be studied within the classical network-theory formalism.

### The basic processes

Biotic interactions themselves interact (Golubski and Abrams 2011); in other words, interactions are contingent on the occurrence of species other than those interacting. Because the outcome of an interaction ultimately affects local abundances (over ecological time scales) and population trait structure (over evolutionary time scales), all interactions happening within a community will impact one another. This does not actually mean pairwise approaches are bound to fail, but it does clamor for a larger scale approach that accounts for indirect effects. The occurrence or absence of a biotic interaction can either affect either the realization of other interactions (thus affecting the “interaction” component of network *β*-diversity) or the presence of other species. There are several well-documented examples of one interaction allowing new interactions to happen, *e.g.* opportunistic pathogens have a greater success of infection in hosts which are already immunocompromised by previous infections, (Olivier 2012), or conversely preventing them, *e.g.* a resident symbiont decreases the infection probability of a new pathogen (Heil and McKey 2003, Koch and Schmid-Hempel 2011). In both cases, the driver of interaction turnover is the patchiness of species distribution; the species acting as a “modifier” of the probability of interaction is only partially present throughout the range of the other two species, thus creating a mosaic of different interaction configurations. Variation in interaction structure can happen through both cascading and environmental effects: Singer et al. (2004) show that caterpillars change the proportion of different plant species in their diet when parasitized in order to favor low quality items and load themselves with chemical compounds which are toxic for their parasitoids. However, low quality food results in birds having a greater impact on caterpillar populations (Singer et al. 2012). It is noteworthy that in this example, the existence of a parasitic interaction will affect both the strength, and impact, of other interactions. In terms of their effects on network *β*-diversity, indirect effects are thus likely to act on components of dissimilarity. A common feature of the examples mentioned here is that pinpointing the exact mechanism through which interactions affect each other often requires a good working knowledge of the system’s natural history.

### Benefits for network analysis

As discussed in previous sections, improved understanding of why and where species interact should also provide a mechanistic understanding of observed species co-occurrences. However, the presence of species is also regulated by indirect interactions. Recent experimental results showed that some predator species can only be maintained if another predator species is present, since the latter regulates a competitively superior prey and allows for prey coexistence (Sanders and Veen 2012). These effects involving several species and several types of interactions across trophic levels are complex (and for this reason, have been deemed unpredictable in the past, Tack et al. (2011)), and can only be understood by comparing communities in which different species are present/absent. Looking at figure 1, it is also clear that the probability of having an interaction between species *i* and *j* (P(**L**_*ij*_)) is ultimately constrained by the probability that individuals of species *i* and *j* will meet assuming random movement, *i.e.* P(*i*∩*j*). Thus, the existence of any ecological interaction will be contingent upon *other* ecological interactions driving local co-occurrence (Araújo et al. 2011). Based on this argument, ecological networks cannot be limited to a collection of pairwise interactions. Our view of them needs be updated to account for the importance of the context surrounding these interactions (Box 2). From a biogeographic standpoint, it requires us to develop a theory based on interaction co-occurrence in addition to the current knowledge encompassing only species co-occurrence. Araújo et al. (2011) and Allesina and Levine (2011) introduced the idea that competitive interactions can leave a signal in species co-occurrence network. A direct consequence of this result is that, for example, trophic interactions are constrained by species’ competitive outcomes *before* they are ever constrained by *e.g.* predation-related traits. In order to fully understand interactions and their indirect effects, however, there is a need to develop new conceptual tools to *represent* effects that interactions have on one another. In a graph theoretical perspective, this would amount to establishing edges between pairs of edges, a task for which there is limited conceptual or methodological background.

## Conclusions

Overall, we argue here that the notion of “species interaction networks” shifts our focus away from the level of organization at which most of the relevant biogeographic processes happen — populations. In order to make reliable predictions about the structure of networks, we need to understand what triggers variability of ecological interactions. In this contribution, we have outlined that there are several direct (abundance-based and trait-based) and indirect (biotic modifiers, indirect effects of co-occurrence) effects to account for. We expect that the relative importance of each of these factors and how precisely they affect the probability of establishing an interaction are likely system-specific; nonetheless, we have proposed a unified conceptual approach to understand them better.

At the moment, the field of community ecology is severely data-limited to tackle this perspective. Despite the existence of several spatially- or temporally-replicated datasets (*e.g.* Schleuning et al. 2011 2012 Menke et al. 2012), it is rare that all relevant information has been measured independently. It was recently concluded, however, that even a reasonably small subset of data can be enough to draw inferences at larger scales (Gravel et al. 2013). Paradoxically, as tempting as it may be to sample a network in its entirety, the goal of establishing global predictions might be better furthered by extremely-detailed characterization of a more modest number of interactions (Rodriguez-Cabal et al. 2013). Assuming that there are in-deed statistical invariants in the rules governing interactions, this information will allow us to make verifiable predictions on the structure of the networks. Better still, this approach has the potential to substantially strengthen our understanding of the interplay between traits and neutral effects. Blüthgen et al. (2008) claim that the impact of traits distribution on network structure can be inferred simply by removing the impact of neutrality (population densities), based on the idea that many rare links were instances of sampling artifacts. As illustrated here (e.g, Box 2), their approach is of limited generality, as the abundance of a species itself can be directly driven by factors such as trait-environment matching. In addition, there are virtually no datasets that follow a collection of interacting species through both space *and* time in a replicated fashion. This type of data, although exceedingly tedious to collect, would provide important indications of which mechanisms should be explored to improve our understanding the variability of species interactions.

Assuming that suitable and accessible empirical data will inevitably accumulate in the coming years, these approaches will rapidly expand our ability to predict the re-wiring of networks under environmental change. There are two broad mechanisms linking network structure to environmental change: changes in population sizes due to modification of demographic parameters, and plastic or adaptive responses resulting in shifted or disrupted trait distributions. The framework proposed in Box 1 predicts interaction probabilities under different scenarios. Ultimately, being explicit about the trait-abundance-interaction feedback will provide a better understanding of short-term and long-term dynamics of interaction networks. We illustrate this in Fig. 3. The notion that population sizes have direct effects on the existence of an interaction stands opposed to classical consumer-resource theory, which is one of the bases of network analysis. Considering this an opposition, however, is erroneous. Consumer-resource theory considers a strong effect of abundance on the intensity of interactions (Box 2), and itself is a source of (quantitative) variation. Furthermore, these models are entirely determined by variations in population sizes in the limiting case where the coefficient of interactions are similar. As such, any approach seeking to understand the variation of interactions over space ought to consider that local densities are not only a consequence, but also a predictor, of the probability of observing an interaction. The same reasoning can be held for local trait distributions, although over micro-evolutionary time-scales. While trait values determine whether two species are able to interact, they will be modified by the selective effect of species interacting. Therefore, conceptualizing interactions as the outcome of a probabilistic process regulated by local factors, as opposed to a constant, offers the unprecedented opportunity to investigate feedbacks between different time scales. This is especially important since all of the mechanisms mentioned above are also likely to change rapidly over spatial scales. The situation in which the phenologies of populations are synchronized locally but not regionally (as shown by Singer and McBride 2012) is an excellent example of when we must integrate these mechanisms into our interpretation of spatial and temporal dynamics.

**Figure 3:**
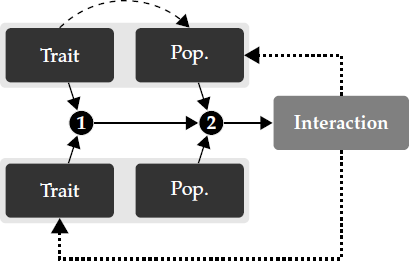
The approach we propose (that populations can interact at the conditions that 1 their trait allow it and 2 they are locally abundant enough for some of their individuals to meet by chance) requires an increased focus on population-level processes. A compelling argument that supports working at this level of organisation is that eco-evolutionary feedbacks are explicit. All of the components of interaction variability we described are potentially related, either through variations of population sizes due to the interaction itself, or due to selection arising from these variations in population size. In addition, some traits involved in the existence of the interaction may also affect local population abundance.

Over the past decade, many insights have been gained by looking at the turnover of different facets of biodiversity (taxonomic, functional, and phylogenetic) through space (Devictor et al. 2010b, Meynard et al. 2011). Here, we propose that there is another oft-neglected side of biodiversity: species interactions. The perspective we bring forth allows us to unify these dimensions and offers us the opportunity to describe the biogeographic structure of all components of community and ecosystem structure simultaneously.

## Acknowledgements

We thank Michael C. Singer and one anonymous referee for insightful comments on this manuscript. TP, DBS, and DG received financial support from the Canadian Institute of Ecology and Evolution (*Continental Scale Variation of Ecological Networks* thematic working group). TP was funded by a FRQNT-MELS PBEE post-doctoral scholarship. DBS was funded by a Marsden Fund Fast-Start grant (UOC-1101) administered by the Royal Society of New Zealand.

## References

Allesina, S. and Levine, J. 2011. A competitive network theory of species diversity. - Proceedings of the National Academy of Sciences of the United States of America 108: 5638.

Allesina, S. and Tang, S. 2012. Stability criteria for complex ecosystems. - Nature 483: 205–208.

Angilletta, M. J. et al. 2004. Temperature, Growth Rate, and Body Size in Ectotherms: Fitting Pieces of a Life-History Puzzle. - Integrative and Comparative Biology 44: 498–509.

Araújo, M. B. et al. 2011. Using species co-occurrence networks to assess the impacts of climate change. - Ecography 34: 897–908.

Baiser, B. et al. 2012. Geographic variation in network structure of a nearctic aquatic food web.- Global Ecology and Biogeography 21: 579–591.

Baskerville, E. B. et al. 2011. Spatial Guilds in the Serengeti Food Web Revealed by a Bayesian Group Model (LA Meyers, Ed.). - PLoS Computational Biology 7: e1002321.

Berlow, E. L. et al. 2009. Simple prediction of interaction strengths in complex food webs. - Proceedings of the National Academy of Sciences of the United States of America 106: 187–91.

Blüthgen, N. et al. 2006. Measuring specialization in species interaction networks. - BMC ecology 6: 9.

Blüthgen, N. et al. 2008. What do interaction network metrics tell us about specialization and biological traits? - Ecology 89: 3387–99.

Bolnick, D. I. et al. 2003. The ecology of individuals: incidence and implications of individual specialization. - The American Naturalist 161: 1–28.

Bolnick, D. I. et al. 2011. Why intraspecific trait variation matters in community ecology. - Trends in Ecology and Evolution 26: 183–192.

Brose, U. et al. 2006. Allometric scaling enhances stability in complex food webs. - Ecology Letters 9: 1228–1236.

Burdon, J. J. and Thrall, P. H. 2009. Coevolution of plants and their pathogens in natural habitats. - Science 324: 755.

Canard, E. et al. 2012. Emergence of Structural Patterns in Neutral Trophic Networks. - PLoS One 7: e38295.

Canard, E. F. et al. 2014. Empirical Evaluation of Neutral Interactions in Host-Parasite Networks. - The American Naturalist 183: 468–479.

Choh, Y. et al. 2012. Predator-prey role reversals, juvenile experience and adult antipredator behaviour. - Scientific Reports in press.

Dáttilo, W. et al. 2013. Spatial structure of ant–plant mutualistic networks. - Oikos: no–no.

Devictor, V. et al. 2010a. Defining and measuring ecological specialization. - Journal of Applied Ecology 47: 15–25.

Devictor, V. et al. 2010b. Spatial mismatch and congruence between taxonomic, phylogenetic and functional diversity: the need for integrative conservation strategies in a changing world.- Ecology Letters 13: 1030–1040.

Devictor, V. et al. 2012. Differences in the climatic debts of birds and butterflies at a continental scale. - Nature Climate Change 2: 121–124.

Diniz-Filho, J. A. F. and Bini, L. M. 2008. Macroecology, global change and the shadow of forgotten ancestors. - Global Ecology and Biogeography 17: 11–17.

Duffy, J. E. et al. 2007. The functional role of biodiversity in ecosystems: incorporating trophic complexity. - Ecology Letters 10: 522–538.

Dunne, J. A. 2006. The Network Structure of Food Webs. - In: Dunne, J. A. and Pascual, M. (eds), Ecological networks: Linking structure and dynamics. Oxford University Press, ppp. 27–86.

Dunne, J. A. et al. 2002. Network structure and biodiversity loss in food webs: robustness increases with connectance. - Ecology Letters 5: 558–567.

Eklof, A. et al. 2011. Relevance of evolutionary history for food web structure. - Proceedings of the Royal Society B: Biological Sciences 279: 1588–1596.

Estes, J. A. et al. 2011. Trophic Downgrading of Planet Earth. - Science 333: 301–306.

Eveleigh, E. S. et al. 2007. Fluctuations in density of an outbreak species drive diversity cascades in food webs. - Proceedings of the National Academy of Sciences of the United States of America 104: 16976–16981.

Gandon, S. et al. 2008. Host-parasite coevolution and patterns of adaptation across time and space. - Journal of Evolutionary Biology 21: 1861–1866.

Gilman, S. E. et al. 2010. A framework for community interactions under climate change. - Trends in Ecology and Evolution 25: 325–331.

Golubski, A. J. and Abrams, P. A. 2011. Modifying modifiers: what happens when interspecific interactions interact? - Journal of Animal Ecology 80: 1097–1108.

Gomulkiewicz, R. et al. 2000. Hot spots, cold spots, and the geographic mosaic theory of coevolution. - The American Naturalist 156: 156–174.

Gravel, D. et al. 2011. Trophic theory of island biogeography. - Ecology Letters 14: 1010–1016.

Gravel, D. et al. 2013. Inferring food web structure from predator-prey body size relationships.- Methods in Ecology and Evolution in press.

Havens, K. 1992. Scale and structure in natural food webs. - Science 257: 1107–1109.

Heil, M. and McKey, D. 2003. Protective ant-plant interactions as model systems in ecological and evolutionary research. - Annual Review of Ecology, Evolution, and Systematics 34: 425–553.

Herrera, C. M. et al. 2002. Interaction of pollinators and herbivores on plant fitness suggests a pathway for correlated evolution of mutualism-and antagonism-related traits. - Proceedings of the National Academy of Sciences 99: 16823–16828.

Holt, R. D. and Kotler, B. P. 1987. Short-Term Apparent Competition. - The American Naturalist 130: 412–430.

Hubbell, S. P. 2001. The Unified Neutral Theory of Biodiversity and Biogeography. - Princeton University Press.

Ings, T. C. et al. 2009. Ecological networks–beyond food webs. - Journal of Animal Ecology 78: 253–269.

Janzen, D. H. and Martin, P. S. 1982. Neotropical anachronisms: the fruits the gomphotheres ate. - Science 215: 19–27.

Kéfi, S. et al. 2012. More than a meal… integrating non-feeding interactions into food webs. - Ecology Letters 15: 291–300.

Koch, H. and Schmid-Hempel, P. 2011. Socially transmitted gut microbiota protect bumble bees against an intestinal parasite. - PNAS: 1110474108.

Koen-Alonso, M. 2007. A process-oriented approach to the multispecies functional response. - In: From energetics to ecosystems: the dynamics and structure of ecological systems. Springer, ppp. 1–36.

Krishna, A. et al. 2008. A neutral-niche theory of nestedness in mutualistic networks. - Oikos 117: 1609–1618.

May, R. M. 1973. Stability in randomly fluctuating versus deterministic environments. - American Naturalist: 621–650.

Memmott, J. et al. 2004. Tolerance of pollination networks to species extinctions. - Proceedings of the Royal Society B: Biological Sciences 271: 2605–2611.

Menke, S. et al. 2012. Plant-frugivore networks are less specialized and more robust at forest- farmland edges than in the interior of a tropical forest. - Oikos 121: 1553–1566.

Meynard, C. N. et al. 2011. Beyond taxonomic diversity patterns: how do \$\$, \$\$ and \$\$ components of bird functional and phylogenetic diversity respond to environmental gradients across France? - Global Ecology and Biogeography 20: 893–903.

Nuismer, S. L. et al. 2003. Coevolution between hosts and parasites with partially overlapping geographic ranges. - Journal of Evolutionary Biology 16: 1337–1345.

Ohba, S.-y. 2011. Field observation of predation on a turtle by a giant water bug. - Entomological Science 14: 364–365.

Olesen, J. M. et al. 2011. Missing and forbidden links in mutualistic networks. - Proceedings. Biological sciences / The Royal Society 278: 725–32.

Olivier, L. 2012. Are Opportunistic Pathogens Able to Sense the Weakness of Host through Specific Detection of Human Hormone? - Journal of Bacteriology & Parasitology in press.

Parmesan, C. 2007. Influences of species, latitudes and methodologies on estimates of phenological response to global warming. - Global Change Biology 13: 1860–1872.

Piechnik, D. A. et al. 2008. Food-web assembly during a classic biogeographic study: species’“trophic breadth” corresponds to colonization order. - Oikos 117: 665–674.

Poisot, T. et al. 2011. Resource availability affects the structure of a natural bacteria-bacteriophage community. - Biology Letters 7: 201–204.

Poisot, T. et al. 2012. The dissimilarity of species interaction networks. - Ecology Letters 15: 1353–1361.

Price, P. W. 2003. Macroevolutionary Theory on Macroecological Patterns. - Cambridge University Press.

Rezende, E. L. et al. 2007. Non-random coextinctions in phylogenetically structured mutualistic networks. - Nature 448: 925–8.

Rodriguez-Cabal, M. A. et al. 2013. Node-by-node disassembly of a mutualistic interaction web driven by species introductions. - Proceedings of the National Academy of Sciences 110: 16503–16507.

Rohr, R. P. et al. 2010. Modeling food webs: exploring unexplained structure using latent traits.- The American naturalist 176: 170–7.

Saavedra, S. et al. 2011. Strong contributors to network persistence are the most vulnerable to extinction. - Nature 478: 233–235.

Sanders, D. and Veen, F. J. F. van 2012. Indirect commensalism promotes persistence of secondary consumer species. - Biology Letters: 960–963.

Schleuning, M. et al. 2011. Specialization and interaction strength in a tropical plant-frugivore network differ among forest strata. - Ecology 92: 26–36.

Schleuning, M. et al. 2012. Specialization of Mutualistic Interaction Networks Decreases to- ward Tropical Latitudes. - Current biology 22: 1925–31.

Singer, M. C. and McBride, C. S. 2012. Geographic mosaics of species’ association: a definition and an example driven by plant/insect phenological synchrony. - Ecology: 120613103411007.

Singer, M. S. et al. 2004. Disentangling food quality from resistance against parasitoids: diet choice by a generalist caterpillar. - The American Naturalist 164: 423–429.

Singer, M. S. et al. 2012. Tritrophic interactions at a community level: effects of host plant species quality on bird predation of caterpillars. - The American naturalist 179: 363–74.

Stouffer, D. B. et al. 2005. Quantitative patterns in the structure of model and empirical food webs. - Ecology 86: 1301–1311.

Stouffer, D. B. et al. 2011. The role of body mass in diet contiguity and food-web structure. - Journal of Animal Ecology: no–no.

Stouffer, D. B. et al. 2012. Evolutionary Conservation of Species’ Roles in Food Webs. - Science 335: 1489–1492.

Tack, A. J. M. et al. 2011. Can we predict indirect interactions from quantitative food webs?–an experimental approach. - The Journal of animal ecology 80: 108–118.

Thompson, J. N. 2005. The Geographic Mosaic of Coevolution. - University Of Chicago Press.

Thuiller, W. et al. 2013. A road map for integrating eco-evolutionary processes into biodiversity models. - Ecology Letters 16: 94–105.

Tylianakis, J. M. et al. 2007. Habitat modification alters the structure of tropical host–parasitoid food webs. - Nature 445: 202–205.

Vazquez, D. P. et al. 2005. Species abundance and the distribution of specialization in host-parasite interaction networks. - Journal of Animal Ecology 74: 946–955.

Vázquez, D. P. et al. 2007. Species abundance and asymmetric interaction strength in ecological networks. - Oikos 116: 1120–1127.

Violle, C. et al. 2012. The return of the variance: intraspecific variability in community ecology.- Trends in Ecology and Evolution 27: 244–252.

Williams, R. and Martinez, N. 2000. Simple rules yield complex food webs. - Nature 404: 180–183.

Woodward, G. et al. 2010. Ecological networks in a changing climate. - Advances in Ecological Research 42: 71–138.

Woodward, G. et al. 2012. Climate change impacts in multispecies systems: drought alters food web size structure in a field experiment. - Philosophical Transactions of the Royal Society B: Biological Sciences 367: 2990–2997.

Wootton, J. T. 2005. Field parameterization and experimental test of the neutral theory of biodiversity. - Nature 433: 309–12.

Yeakel, J. D. et al. 2012. Probabilistic patterns of interaction: the effects of link-strength variability on food web structure. - Journal of The Royal Society Interface: rsif.2012.0481.

